# Pyro-Velocity: Probabilistic RNA Velocity inference from single-cell data

**DOI:** 10.1101/2022.09.12.507691

**Authors:** Qian Qin, Eli Bingham, Gioele La Manno, David M. Langenau, Luca Pinello

## Abstract

Single-cell RNA Velocity has dramatically advanced our ability to model cellular differentiation and cell fate decisions. However, current preprocessing choices and model assumptions often lead to errors in assigning developmental trajectories. Here, we develop, Pyro-Velocity, a Bayesian, generative, and multivariate RNA Velocity model to estimate the uncertainty of cell future states. This approach models raw sequencing counts with the synchronized cell time across all expressed genes to provide quantifiable and improved information on cell fate choices and developmental trajectory dynamics.

## Main text

RNA Velocity is a powerful computational framework to estimate the time derivative of gene expression^1–4^, model transcriptional dynamics, and predict cell fate from single-cell RNA sequencing (RNA-seq) data. The framework has been used to study developmental cell lineage trajectories over a broad spectrum of biological processes, such as haematopoiesis^3^, pancreas development^5^, and neural progenitors differentiation^4^. Yet, recent state-of-the-art RNA Velocity approaches such as *velocyto* and *scVelo*^1^ have several limitations. First, they may predict developmental trajectories that do not exist in nature, as it has been shown in the analysis of fully mature blood cells^6,7^. Second, data preprocessing steps (e.g., selection of top variable genes, dimensionality reduction) affect velocity estimates that are difficult to characterize and might lead to artefactual predictions. In addition, non-linear spliced and unspliced imputation procedures could distort phase portrait geometry and recovered trajectories (**Fig. 1a**). Third, most RNA velocity algorithms perform estimates independently for each gene, and therefore cannot exploit the natural interdependence of gene expression. Fourth, a unified description of the temporal relation between cells can only be assigned based on postprocessing steps that aggregate heuristically the gene-wise fits into a shared latent time (**Fig. 1a**). Finally, current RNA velocity frameworks only output point estimates of the velocity vector field and do not provide uncertainty estimation. Importantly, the absence of statistical confidence estimates increases false discoveries as it leaves users without a way to evaluate the robustness of the methods’ outputs. To overcome these limitations, we developed a probabilistic generative model, named *Pyro-Velocity*, that estimates RNA dynamics and infers cell trajectories using Pyro^8^, a robust and efficient probabilistic programming language.

**Figure 1.**
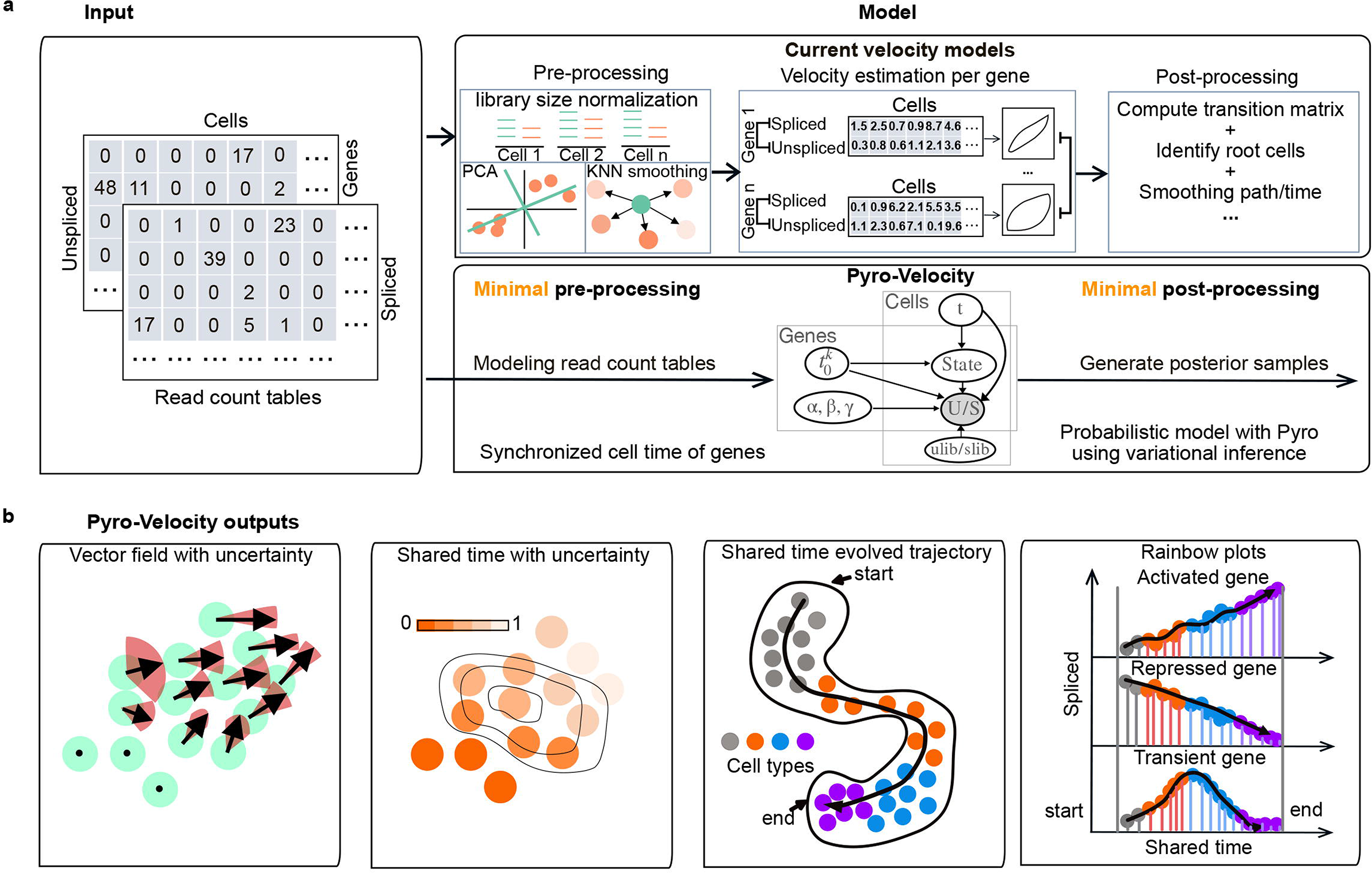
Pyro-Velocity is a fully generative Bayesian method with uncertainty estimation of velocity vector fields and shared latent time, based on raw counts and without ad-hoc preprocessing steps. **a.** Schematic comparing current RNA Velocity and Pyro-Velocity workflows. Both methods take in input spliced and unspliced read count tables and output RNA velocity vector fields and latent (shared) time. Previous methods require ad-hoc preprocessing steps, including the arbitrary selection of top variable genes, number of principal components, and kNN-pooling to impute both spliced and unspliced expression within the PCA-reduced spliced expression space. In addition, manually tuned ad-hoc postprocessing steps are used to compute the final vector field and latent time based on aggregation and smoothing of the independent velocity estimates for each gene which can dramatically change the velocity estimates. In contrast, Pyro-Velocity is a fully generative and probabilistic model that uses raw read counts. Our model requires minimal preprocessing and postprocessing steps and outputs a common *shared* time based on all the genes without additional aggregation steps. It also provides uncertainty estimation based on posterior samples. **b.** The main outputs of the Pyro-Velocity framework. Visualization of single gene expression across inferred developmental cell order using rainbow plots (right panel).

Pyro-Velocity recasts the velocity estimation problem into a latent variable posterior probability inference (**Methods**). The proposed model is generative and fully Bayesian, with the different parameters considered as latent random variables. Each parameter is assigned a prior distribution, and the model is then conditioned on the raw data (both spliced and unspliced molecule raw counts) to estimate posterior probability distributions and to enable uncertainty estimation of velocity. Central to the Pyro-Velocity model is a *shared latent time* (*t*): a global latent variable placing each cell at a particular instant of the modeled dynamical process. From this latent time, the expectation of spliced and unspliced RNA levels for each gene can be computed knowing gene-specific kinetics parameters (transcriptional (*α*), splicing (*β*), and degradation rates (*γ*) and two gene-specific switching times 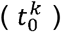, and gene-specific transcriptional state *k*) and integrating the RNA velocity differential equations analogously to what proposed in the dynamical RNA velocity model by Bergen et al.^4,9^. However, we innovate the structure and formulation of the latent time given that our model is set to jointly fit the entire raw data matrix, not individual genes. We approach the task of estimating the posterior distribution by restricting the search within a family of a variational distribution approximating the true posterior, and taking advantage of the automatic differentiation variational inference (ADVI) capabilities of Pyro^8,10^ (**Fig. 1a**, **Methods**). Notably, our generative Bayesian model enables to sample from the posterior distributions of the latent and dependent variables, allowing uncertainty estimation of phase portraits, high dimensional velocity vector field, and shared latent time. Around this model, we built several visualizations to assess uncertainty and functions to recover putative cell fate priming genes and to investigate how the denoised gene expression recapitulates known cell-type annotations and their developmental order (**Fig. 1b, Methods**). Although several RNA velocity methods^11–15^ have been recently proposed, and many of them have not been peer-reviewed, here we showcase the advantages and innovations of Pyro-Velocity by comparing it with the current state-of-the-art dynamical RNA velocity model proposed by Bergen et al.^4^, and implemented in scVelo.

One major biological hurdle for current formulations of RNA Velocity models has been miscalling cell trajectories in fully differentiated cell populations that lack dynamic cell state transitions. One typical example being human peripheral blood mononuclear cells (PBMCs)^6^. In this dataset, 65,877 cells were analyzed using the 10x platform that comprised 11 fully differentiated, mature immune cell types. This dataset lacked stem and progenitor cells or other signatures of an undergoing dynamical differentiation; thus, no consistent velocity flow should be detected. We reanalyzed this dataset using scVelo and Pyro-Velocity using a common set of genes for a fair comparison. Using scVelo, we reproduced the original results showing incorrect cell trajectory assignments (**Fig. 2a**)^7^. ScVelo uncovered an errant, strong linear trajectory across a subset of mature cell types^6^ that is exasperated by the inability to confidently assess lineage directionality or a baseline to assess significance. On the contrary, Pyro-Velocity failed to detect high-confidence trajectories in the mature blood cell states, consistent with what is known about the biology underlying these cells (**Fig. 2a,** panel 2 vs. panel 3). Importantly, the Bayesian nature of Pyro-Velocity enabled the generation of uncertainty estimates that can be easily visualized as likely vectors for individual cells or averaged across cells on a constantly spaced grid on the cell embedding (*averaged vector field*).

**Figure 2.**
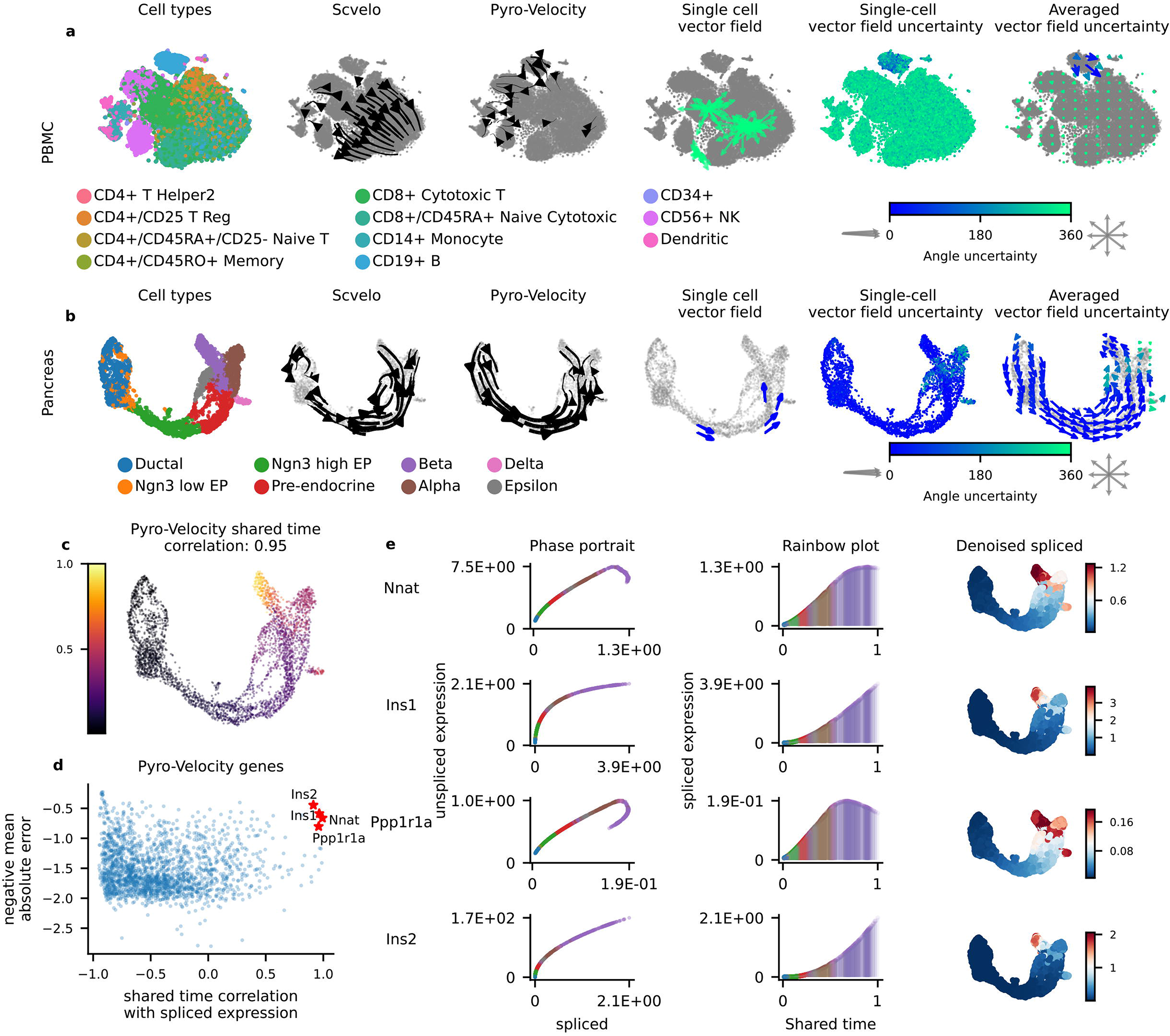
Pyro-Velocity outperforms other RNA-velocity methods and accurately predicts cell lineage trajectories. **a,b.** RNA Velocity model comparisons from fully mature peripheral blood mononuclear cells (PBMCs, t-SNE embedded plots, a) or E15.5 mouse pancreas (UMAP embedded plots, b-c). From left to right in panels a and b: 1. Cells colored by cell type; 2. Stream plot of velocity vector field defined by scVelo; 3. Stream plot of the posterior mean vector field from Pyro-Velocity; 4. Pyro-Velocity visualization of cell dynamics for 3 arbitrarily selected single cells; 5. *Single-cell vector field uncertainty* where each cell is colored based on the scaled angular standard deviation to visualize uncertainty based on n=30 iterated, posterior samples depicted as arrows noted throughout our analysis; 6. *Averaged vector field uncertainty* where each vector is colored by the averaged vector field uncertainty. The scaled angular standard deviation range is [0 360], where 0 corresponds to the highest confidence for a single direction, 360 corresponds to the highest uncertainty, and no directionality preference based on 30 posterior samples (4-6). Vectors were projected into the UMAP space using the scVelo API based on cosine similarity-derived transition matrix. **c.** UMAP embedded cells colored by the average cell shared time across 30 posterior samples. The Spearman’s correlation between Pyro-Velocity and Cytotrace is provided. **d.** Pyro-Velocity automatic gene selection and ranking for predicting important genes involved in pancreas development. **e.** Phase portraits of posterior samples average spliced and unspliced gene expression (left), rainbow plots showing denoised spliced gene expression (middle), and gene expression within UMAP embedded space. Single dots correspond to individual cells (left, right) or vertical lines (middle). Cell type annotation coloring is the same as in panel b.

Pyro-Velocity was also able to define well-known developmental cell hierarchies in a scRNA-seq dataset of E15.5 mouse pancreas (**Fig. 2b-e**, n=3,696 cells comprising 8 cell types (**Fig. 2b,** panel 1)), identifying cell trajectories originating from ductal progenitor cells and culminated in the production of mature Alpha, Beta, Delta, and Epsilon cells^5^. Though the scVelo and Pyro-Velocity models have qualitatively comparable vector fields, the results generated by scVelo were greatly influenced by gene selection, neighboring cell number for gene expression smoothing, and cannot provide confidence assessment of lineage directionality (**Supplementary Fig.2a**). In addition, Pyro-Velocity uncertainty visualizations uncovered that differentiated cells had a larger vector field uncertainty than stem and progenitor cells, as would be expected for terminally differentiated cell states where progression and lineage trajectories would not be anticipated. Pyro-Velocity also recovered a developmental order in close agreement with what was estimated by Cytotrace (Spearman’s r=0.95, p-value<1E-10 between latent time and developmental order)^16^ - a state-of-the-art orthogonal method used to predict cell differentiation based on the number of expressed genes per cell.

We next used Pyro-Velocity to unbiasedly uncover potential cell fate determinant markers genes within the E15.5 mouse pancreas. To this end, we first selected the top 300 genes with the best goodness of fit (negative mean absolute errors across cells). Next, we ranked them based on the positive (or negative) correlation between inferred shared time and corresponding denoised spliced expression per cell (**Fig. 2d, Methods**). This procedure successfully recovered key developmental genes that are activated (or repressed) during endocrinogenesis (**Fig. 2d-e**, **Supplementary Fig. 2**). Among the top positively correlated genes identified by Pyro-Velocity, endocrine Beta cells expressed *insulin 1* and *2* (*Ins1, Ins2*) and *Nnat,* which is a key regulator Beta cell insulin generation^17^ and *Ppp1r1a* which is a marker of Beta cells^18^. The ranking procedure proposed by scVelo failed to identify *Ins1, Ins2, Nnat,* and *Ppp1r1a* as lineage-restricted genes in their top 50 ranked gene list. The inability of scVelo to identify these key developmental genes could be likely explained by the kNN smoothing of read count and filtering steps (**Supplementary Table 1, 2**). Visual inspection of these four top genes by Pyro-Velocity revealed a single activation phase during cell type transitions based on phase portrait curves. We also propose their visualization through “rainbow plots” to illustrate denoised single gene spliced expression across shared time and to investigate the inferred cell order based on known annotations. Further, UMAP embedding of cells colored based on denoised gene expression confirmed the cell fate markers’ smooth and progressive activation (or repression) towards endocrine cells (**Fig. 2e**, **Supplementary Figure 2**).

Ground truth velocity is not a quantity we can easily access. Therefore, no robust and quantitative benchmarking of RNA velocity methods has been proposed. However, recent single-cell assays allow recording lineage relationships and can be used to assess the ability of different methods in recapitulating cell fate progression. Therefore, we next systematically compared scVelo and Pyro-Velocity ability to recover single cell fates in Lineage And RNA RecoverY (LARRY) barcoded hematopoietic cells. LARRY leverages unique lentiviral barcodes inserted into 3’ untranslated region (UTR) of GFP reporter that is integrated as a single copy into parental cells. Then progeny can be clonally traced to follow the descendent cell fates over time. LARRY has been recently applied to *in vitro* differentiation of human blood and accurately recovered expected cell fate and lineage trajectories^19^. This dataset sampled differentiation over the course of 2, 4, and 6 days and contained 49,302 cells that each could be traced with at least one barcode. Because LARRY couples single-cell transcriptomics and direct cell lineage tracing, this data set provides a unique opportunity to benchmark RNA Velocity quantitatively and to provide insights into the correct interpretation of recovered Velocity vector fields and latent cell times. To quantitatively assess the agreement between observed and predicted fates, we considered two metrics: 1. average cosine similarity (*cs*) between vectors from the RNA Velocity vector field and the *clonal progression vector field,* i.e., a vector field derived directly from the lineage data (**Methods**), 2. Spearman’s correlation (*ρ*) between cell latent (shared) time and fate potency scores derived from Cytotrace and Cospar. Cospar, our gold standard, is a state-of-the-art method designed explicitly for predicting fate potency based on LARRY data, while Cytotrace leverages only transcriptomic data. We selected cell state changes associated with monocytes and neutrophils lineages because both lineages have the largest numbers of cells in the scRNA sequencing data set and well-known cell fate trajectories^20^. As expected, the Pyro-Velocity vector field accurately predicted cell state trajectories from hematopoietic progenitor cells into mature monocytes or neutrophils and had high cosine similarity between the inferred vector field and clonal progression vector field assigned by LARRY (monocytes *cs*=0.49, neutrophils *cs*=0.38). High Spearman’s correlation was also seen between shared latent time and Cospar fate potency (monocytes ρ=0.60, neutrophils ρ=0.74) or shared latent time with Cytotrace fate potency (monocytes ρ=0.925, neutrophils ρ=0.925). By contrast, scVelo generated vector fields with reversed order or low correlation (cosine similarity: −0.38, monocytes, cosine similarity: 0.32, neutrophils). Moreover, Cospar fate potency scores were negative (ρ=−0.59, monocytes) or non-significantly correlated (ρ=−0.07, neutrophils). A similar low correlation was observed for scVelo and Cytotrace fate potency scores (monocytes ρ=0.309, neutrophils ρ=0.751) (**Fig. 3a-b, Supplementary Fig. 4c**).

**Figure 3.**
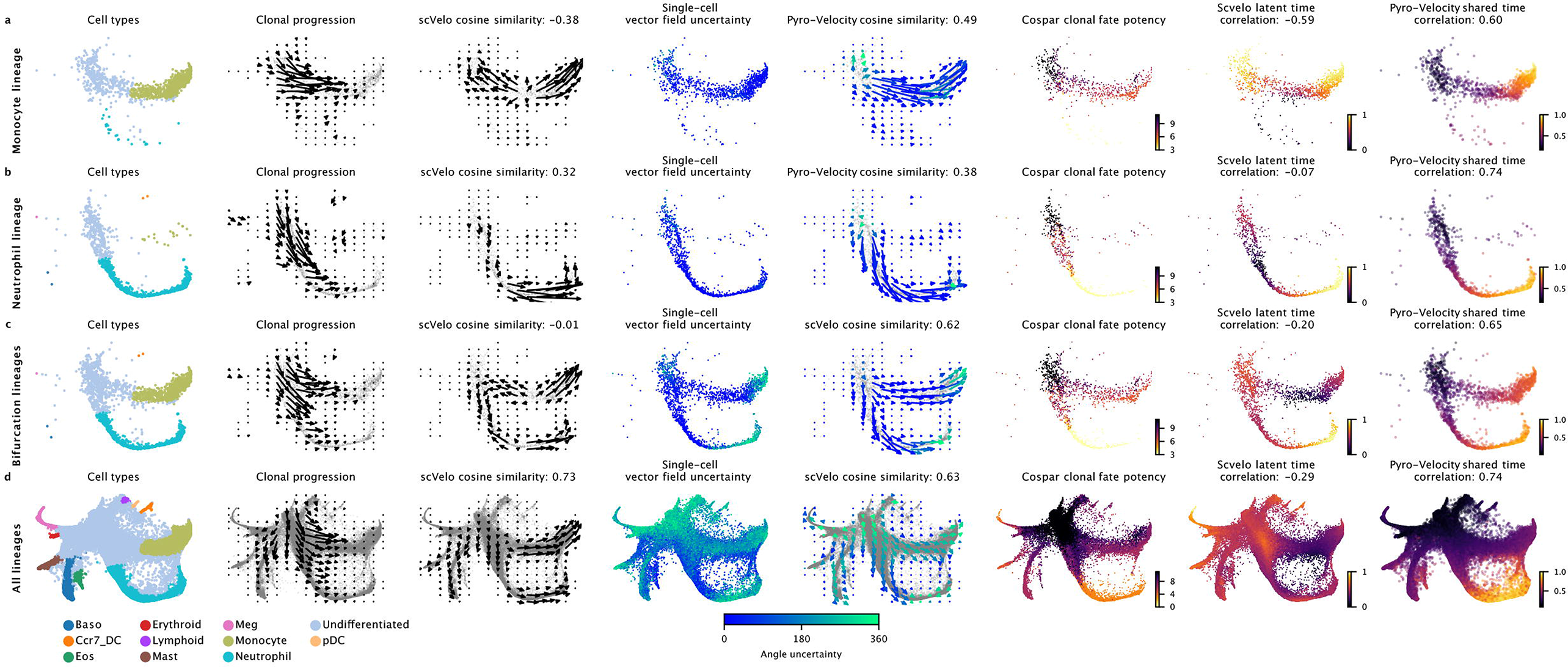
Pyro-Velocity accurately predicts cell fate choices and lineage trajectory in human LARRY barcoded hematopoietic cells. Systematic benchmarking of cell fate prediction methods - predicting uni-lineage cell fates (**a-b**, Model 1), bifurcated lineages (**c**, Model 2), or all lineages together (**d**, Model 2). Panels from left to right show on the same SPRING embedding: 1. Cell types; 2. Clonal progression vector field derived based on the observed clonal barcodes (**Methods**); 3. scVelo recovered vector field; 4. Pyro-Velocity single cell level vector field uncertainty (angular standard deviation from 30 posterior samples); 5. Pyro-Velocity recovered vector field; vectors are colored by the averaged vector field uncertainty; 6. Clonal fate potency score using Cospar^24^; 7. scVelo recovered latent time; 8. Pyro-Velocity inference of posterior mean shared time per cell. Cosine similarity and Spearman’s correlation for the different methods with the clonal progression and fate potency are reported in the titles.

Next, we evaluated the ability of the Pyro-Velocity models to recover bi-fate or multi-fate trajectories by considering clones belonging to monocyte and neutrophil lineages or using the entire dataset. Not surprisingly, both methods showed their limitations, given that they are not designed to model and capture multiple fate trajectories (**Supplementary Fig. 5c-d**). The recovered vector fields and cell latent (shared) time show inconsistent directions for both methods. However, Pyro-Velocity identified higher uncertainty for some of the problematic regions. Furthermore, we discovered that vector field and shared time estimation could be significantly improved by extending the original kinetic model by allowing each gene a time lag for the transcriptional activation and a baseline expression level potentially different than 0 (Pyro-Velocity Model 2, **Methods**) (**Fig. 3c,d, Supplementary Fig. 1b,c**). This extended model recapitulated a vector field consistent with the *clonal progression vector field* for most branches with some spurious predictions within the intermediate cell state between stem cells and Megakaryocytes and most terminal Neutrophils (bi-fate cosine similarity: 0.62, multi-fate cosine similarity: 0.63). However, the proposed uncertainty estimation accurately flagged these subpopulations to warn users of potential false discoveries. Notably, this extended model significantly improved the correlation of cell shared time with the clonal fate potency (bi-fate ρ=0.65, multi-fate ρ=0.74) and also improved the correlation with Cytotrace fate potency (bi-fate ρ=0.931, multi-fate ρ=0.614). Remarkably, the shared time per cell from Pyro-Velocity also outperformed Cytotrace for developmental order prediction (**Supplementary Fig. 4a,c**). Importantly, this extended model retained its ability to accurately identify cell lineage trajectories and shared time for all the other datasets analyzed above **(Supplementary Fig. 3)**. Taken together, our approach using LARRY barcoded datasets established the first quantitative benchmark of RNA velocity-based cell fate trajectory inference and provides an exciting new framework for continued optimization of multi-fate predictions in the future.

In conclusion, Pyro-Velocity provides the first probabilistic, scalable, and end-to-end Bayesian inference framework for RNA Velocity that learns cell dynamics and recovers cell fate from multimodal raw read counts. This approach provides uncertainty estimations of cell fate trajectories and novel visualization approaches to mitigate false discoveries. Pyro-Velocity recovered shared time and vector fields that are recapitulated in true developmental order and outperformed state-of-the-art methods like Cytotrace and scVelo (**Supplementary Fig. 4a**).

## Software availability

We developed our Pyro-Velocity models with Pyro (version 1.6.0) probabilistic programming language and PyTorch (version 1.8.1). Pyro-Velocity source code is available at https://github.com/pinellolab/pyrovelocity. The version of Pyro-Velocity (v0.1.0) used to generate the results presented in this manuscript has been deposited on Zenodo: https://zenodo.org/record/7072876#.Yx-2e-zMLxg.

## Data availability

All the data presented in this manuscript were collected from previously published studies. The fully mature PBMC, pancreas datasets are downloaded and loaded through the scVelo using scvelo.datasets.pancreas(),and scvelo.datasets.pbmc68k(). The LARRY inDrop sequencing data was processed by the LARRY authors using dropEst^20^ and velocyto. This dataset can be accessed at https://figshare.com/articles/dataset/larry_invitro_adata_sub_raw_h5ad/20780344.

## Acknowledgments

This project has been made possible in part by grant number 2022-249212 from the Chan Zuckerberg Initiative DAF, an advised fund of Silicon Valley Community Foundation. L.P. received support from the U.S. NIH (R35 HG010717) and Silicon Valley Community Foundation (2022-249212). D.M.L is supported by R01CA226926, R01 CA215118, R01CA154923, R24 OD031955, CureSearch Acceleration Initiative, and the MGH Scholars Award. We thank the Pinello and Langenau labs members for helpful discussions.

## Author contributions

L.P., Q.Q., and E.B. conceived the work. Q.Q. and E.B created the software. Q.Q. analyzed the data, and prepared the figures and tables. L.P and D.L. supervised the project. L.P., D.L., E.B., G.L.M helped in designing experiments and analyses. All the authors contributed to the writing of the manuscript.

## Competing financial interests statement

L.P. has financial interests in Edilytics, Inc., Excelsior Genomics, and SeQure Dx, Inc. L.P.’s interests were reviewed and are managed by Massachusetts General Hospital and Partners HealthCare in accordance with their conflict of interest policies.

## Methods

### Model formulation

We assume the dynamical gene expression is determined by the RNA splicing process, and infer the unspliced and spliced gene expression level from the differential equations proposed in *velocyto* and scVelo^1,4^:

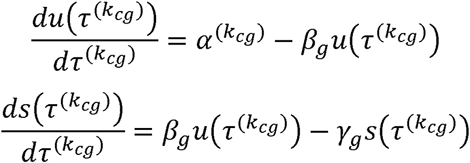

In the equation, the subscript *c* is the cell dimension, *g* is the gene dimension, 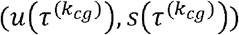 are the unspliced and spliced expression functions given the change of time per cell and gene. τ_*cg*_ represents the displacement of time per cell and gene with 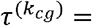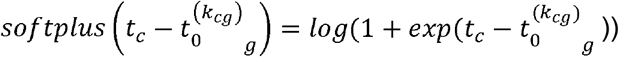, in which *t_c_* is the shared time per cell, 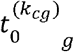 is the gene-specific switching time. Each cell and gene combination has its transcriptional state *k_cg_* ∈ {0,1}, where 0 indicates the activation state and 1 indicates the expression state. Each gene has two switching times for representing activation and repression: 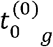 is the first switching time corresponding to when the gene expression starts to be activated, 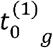 is the second switching time corresponding to when the gene expression starts to be repressed.

We note that *α*^(1)^ is shared for all the genes, while *α*^(0)^_*g*_ is learned independently for each gene. The analytic solution of the differential equations to predict spliced and unspliced gene expression given their parameters is derived by the authors of scVelo^4^ and a theoretic RNA Velocity study^9^.

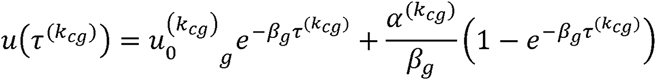

If *β* ≠ *γ*, then

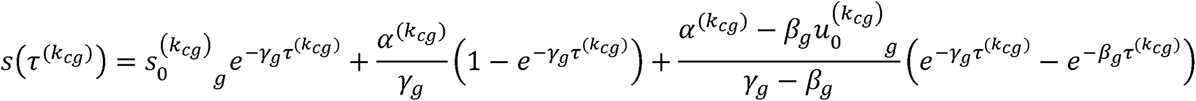

If *β* = *γ*, then

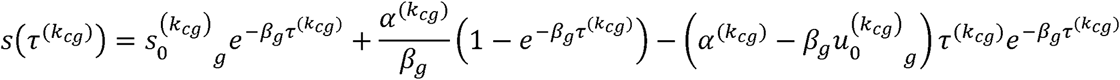

To simplify the above equation, we consider the case when *K_cg_* = 0 and *β_g_* ≠ *γ_g_*, then

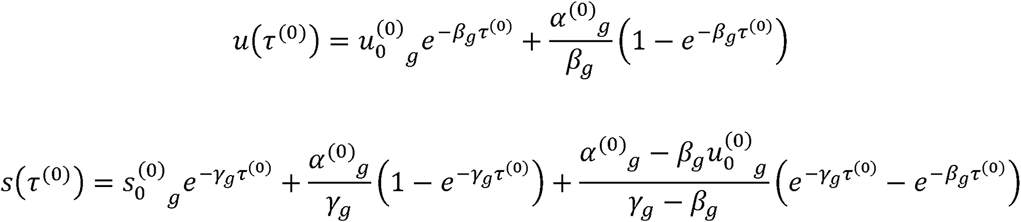

When *k_cg_* =0 and *β_g_* = *γ_g_*, *u*(τ^(0)^) has the same solution, *s*(τ^(0)^) becomes:

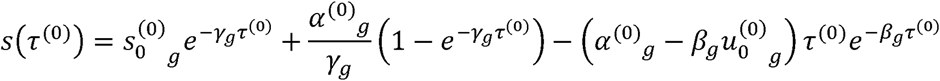

When *k_cg_* =1 and *β_g_* ≠ *γ_g_*, then

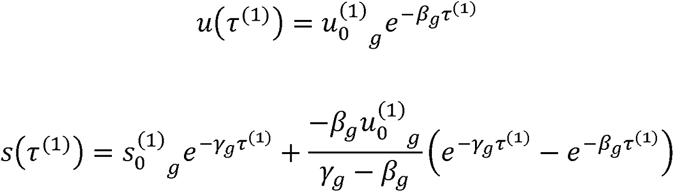

When *k_cg_* =1 and *β_g_* = *γ_g_*, *u*(τ^(1)^) has the same solution, *s*(τ^(1)^) becomes:

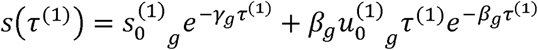

We use these solutions to formulate an end-to-end probabilistic generative model that relates prior distributions on kinetic parameters to a distribution on pairs of observed unspliced and spliced read count matrices:

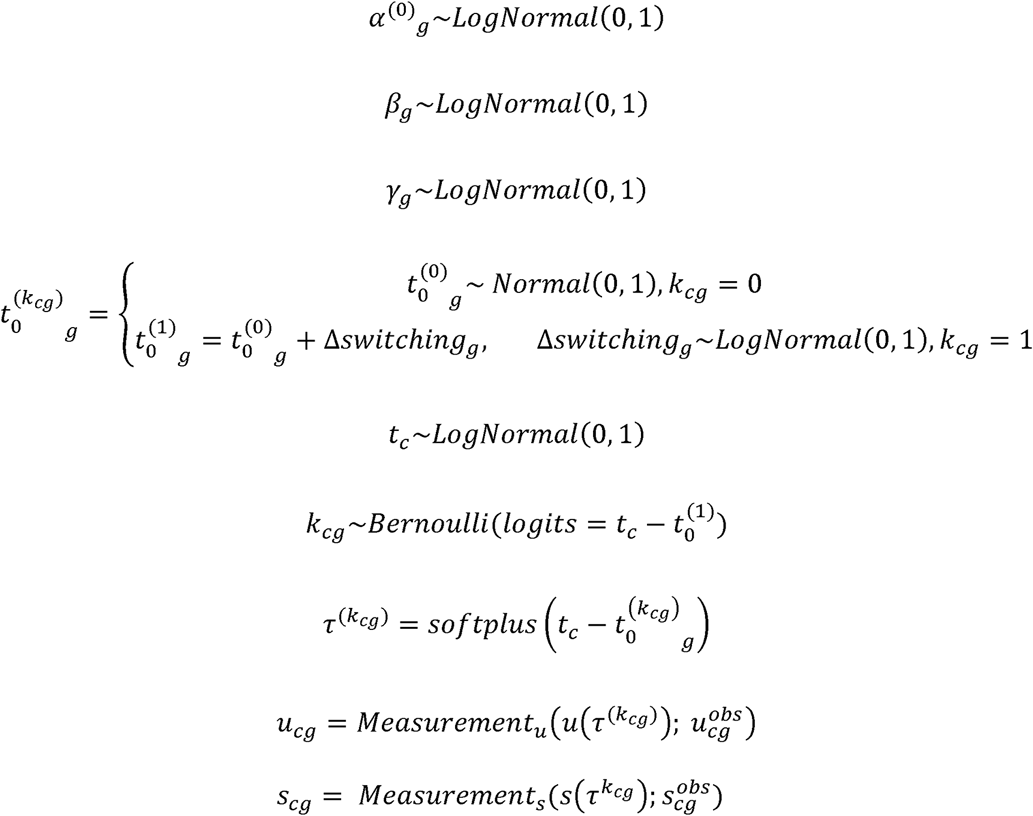

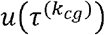 and 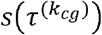 are called the denoised gene expression calculated from the velocity analytic solution input with the kinetics random variables. *u_cg_* and *s_cg_* are the spliced and unspliced read count sampled from the Poisson models. 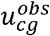 and 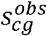 are the observed spliced and unspliced read count tables.

The generative process *Measurement*(.) for observed unspliced read counts given denoised unspliced gene transcript expression level 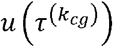 (and identical for observed spliced read counts) models the expected number of observed reads for a given gene in a given cell as the number of transcripts times the ratio of the cell’s total reads to total transcripts:

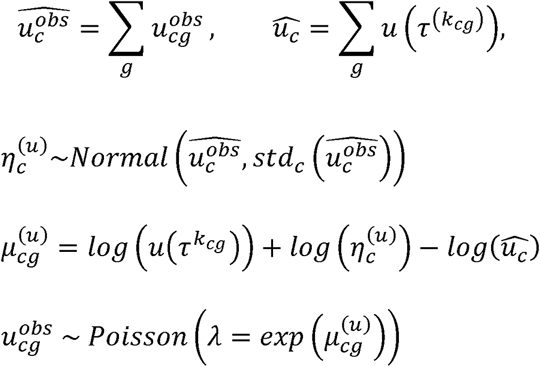

For the first Pyro-Velocity model (Model 1), we constrain the shared time to be strictly larger than 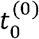 by introducing auxiliary random variables 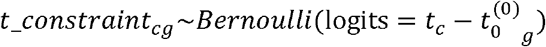 and setting their values to 1, and we set the initial condition per gene to be:

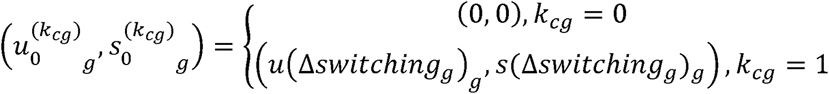

For the extended Pyro-Velocity model (Model 2), we remove the shared time constraint *t_constraint_cg_*, thus allow a time lag per gene that might be caused by delayed gene activation and set the initial condition per gene as random variables that are strictly positive 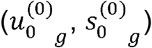, which allow genes having basal expression level before gene activation. Then, we compute the gene expression at the second switching time as 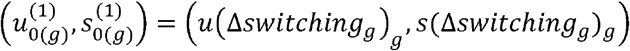 which shares the same initial condition 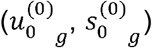 where 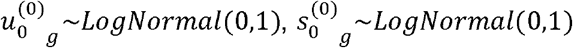.

### Variational inference

Given observations 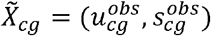, we would like to compute the posterior distribution over the random variables 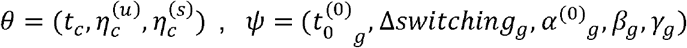, but exact Bayesian inference is intractable in this model. We use Pyro to automatically integrate out the local discrete latent variables *k*, which is defined as the cell and gene transcriptional state (see above), and approximate the posterior over the remaining latent variables using variational inference^8,10^, which converts intractable integrals into optimization problems that can be solved with off-the-shelf tools. In variational inference, we maximize a tractable bound (the evidence lower bound (ELBO)) on the Kullback-Leibler (KL) divergence 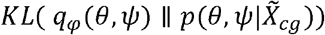 between a parametric family of tractable probability distributions *q* and the intractable true posterior:

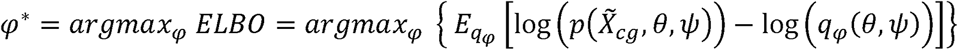

We approximate our model’s posterior distribution 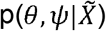 with a tractable family of probability distributions 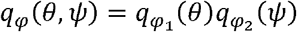, where 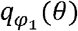 is a product of independent univariate Gaussian distributions with learnable location and scale parameters and 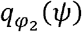 is a multivariate Gaussian distribution with a learnable location parameter and a learnable low-rank covariance matrix.

We solve the resulting stochastic optimization problem using a version of Automatic Differentiation Variational Inference (ADVI)^10^: we obtain gradient estimates by differentiating a Monte Carlo estimate of the ELBO and update the variational parameters *φ* using stochastic gradient ascent. Our implementation and experiments use generic variational families and Monte Carlo ELBO and gradient estimators provided by Pyro^8^, an open source library for probabilistic machine learning, and a version built into Pyro of the adaptive stochastic gradient ascent algorithm Adam augmented with gradient clipping to enhance numerical stability during training.

### Model training

For the pancreas, PBMC, the uni-fate and bi-fate LARRY single-cell data, since the data dimension is relatively small, we run Model 1 and Model 2 with a minimum of 100 epochs and a maximum of 4000 epochs. For each epoch, we input spliced and unspliced read count from all the cells and determine the convergence condition with an early stopping strategy in which the patience is set to 45 and consumes one patience if the minimal ELBO improvement of training data per epoch is lower than 1e-4 of previous loss. The learning rate is set to be 1e-2 with a decay rate of 0.1^1/4000^ per epoch.

For the multi-fate LARRY dataset, since this is a large dataset with over 40K cells, we use mini batches of cells to train Model 1 and Model 2 with a minimum of 100 epochs and a maximum of 1000 epochs. Specifically, the batch size was set to be 4000 cells for both models with an early stopping patience 45 and consume one patience if the minimal ELBO improvement of training data per epoch lower than 1e-3 of previous loss. The learning rate is set to be 1e-2 with a decay rate of 0.1^1/1000^ per epoch.

All models were trained on a machine with an NVIDIA A100 graphic card and the CentOS 7 operating system.

### Posterior prediction

To benchmark Pyro-Velocity performance in predicting cell fate, we generated the posterior samples measurement 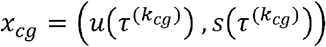 *or x_cg_* = (*u_cg_*, *s_cg_*), *t_c_*, *β_g_*, *γ_g_* from the same single cell RNA-seq using *N* = 30 Monte Carlo samples from the posterior predictive distribution following: *p*(*x_cg_*|*θ,ψ*)*p*(*θ,ψ*|) ≈ *p*(*x_cg_*|*θ,ψ*)*q_φ_*(*θ,ψ*). *x_cg_* can be posterior samples of either denoised gene expression (used for phase portraits and vector field-based trajectory inference) or raw read counts (used for gene selection). Then we calculated the posterior samples of RNA velocity as 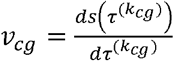 based on posterior samples measurement of denoised gene expression *x_cg_* and *β_g_*,*γ_g_*.

### Prioritization of the cell fate markers

We prioritize the cell fate markers using two metrics: First, the Pearson correlation between each gene’s posterior mean of the denoised spliced expression and posterior mean of the shared time; Second, the negative mean absolute errors of each gene’s observed spliced, and unspliced read counts with posterior predictive samples of spliced and unspliced raw read counts, i.e.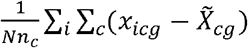, *N* is the posterior sample number that is set to 30, *i* is the posterior sample index, *n_c_* is the cell number. We first select the top 300 genes with the highest negative mean absolute errors and then rank the 300 genes based on the most positively correlated genes and the least negatively correlated genes. We use the same strategy for scVelo to rank the markers by the model likelihood to get the top 300 genes and then prioritize these genes by Pearson’s correlation between scVelo latent time and normalized expression.

### Single-cell data preprocessing

We used Scanpy and scVelo to handle the data input and output; thus, both h5ad and loom files generated by velocyto and kallisto^21^ are supported.

The fully mature PBMC dataset was processed with the same procedure proposed in a recent review paper^7^ (https://scvelo.readthedocs.io/perspectives/Perspectives/). We reproduced this procedure using the scVelo package and raw read counts of the same top three dynamical genes NKG7, IGHM, and GNLY with the best likelihoods.

The pancreas dataset was processed with the scVelo with the following options:

scv.pp.filter_and_normalize(adata, min_shared_counts=30, n_top_genes=2000)
scv.pp.moments(adata, n_pcs=30, n_neighbors=30)

The same top variable genes with raw spliced and unspliced read counts were used as input for the Pyro-Velocity model.

The original LARRY dataset of in vitro Hematopoiesis containing 130,887 cells was first filtered to remove cells without LARRY barcoding. 49,302 cells were recovered after this step with at least one LARRY barcode. For simplicity, we termed this filtered dataset with multiple cell fate (multi-fate) as the full dataset. Based on this dataset, we created two datasets with uni-fate progression toward monocyte or neutrophil based on the lineage LARRY barcodes and time information. Namely, we selected sets of cells with a single LARRY barcode, spanning three time points (day 2, 4, 6), and all the cells from the last time point (day 6) belong to a unique cell type (either monocyte or neutrophil). The two uni-fate datasets were combined to represent the bi-fate LARRY dataset. The multi-fate full dataset was processed using the same options as the pancreas dataset; the rest of the uni-fate and bi-fate datasets were processed using the following parameters:

scv.pp.filter_and_normalize(adata, n_top_genes=2000, min_shared_counts=20)
scv.pp.moments(adata)

### scVelo model

We benchmarked the dynamical RNA velocity model implemented in scVelo (version 0.2.4) for the pancreas and the four LARRY datasets using the same user options :

scvelo.tl.recover_dynamics(adata, n_jobs=30)
scvelo.tl.velocity(adata, mode=‘dynamical’)

Then, we tested a set of user options, including the neighboring cell numbers and the top variable gene numbers, in the pancreas dataset to explore the stability of the scVelo dynamical model. For the fully mature PBMC dataset, we followed the notebook proposed by the original authors (https://scvelo.readthedocs.io/perspectives/Perspectives/), i.e., we used the stochastic RNA velocity model implemented in scVelo with the top three likelihood genes. The latent time from scVelo was computed using their provided function: scvelo.tl.latent_time(adata).

### Trajectories inference by vector fields

#### Velocity vector field

The velocity-based vector fields were generated using the scVelo function as:

scvelo.tl.velocity_graph(adata)
scvelo.tl.velocity_embedding(adata, vkey=‘velocity’, basis=‘embed’)

We used the default options for projecting the vector fields from scVelo models. Unlike scVelo, Pyro-Velocity uses posterior samples (or posterior mean) of the denoised spliced gene expression and posterior samples (or posterior mean) of the velocity estimation for building the transition matrix with cosine similarity. Pyro-Velocity uses the same projection method as scVelo for projecting the transition matrix into the two-dimensional vector field on the user-provided embedding spaces.

#### Clonal progression vector field

No method exists for projecting clonal cell fates from LARRY datasets into a vector field. Thus, we designed a strategy for projecting *clonal progression vector fields:* based on the cells sharing the same LARRY barcode, we considered the vectors connecting the embedding centroid of cells at consecutive time points and then leveraged Gaussian smoothing of nearest centroid directions to represent on a regular grid over the cell manifold the *clonal progression vector field* informed by LARRY barcodes.

#### Cell fate uncertainty estimation

Pyro-Velocity evaluates the cell fate uncertainty by four statistics. The first statistic is the uncertainty of shared time which is evaluated using the standard deviation of posterior samples of *t_c_* for each cell. The second statistic is the angular standard deviation^22^ across posterior samples of the velocity-based vector field. The value of this statistic was mapped linearly to the range [0, 360] for interpretability. The third statistic is the Rayleigh test^23^ for posterior samples of the velocity-based vector fields for each cell, whose p values have been corrected for multiple tests using Benjamini-Hochberg FDR methods; if FDR is less than 0.05, the cell has one statistically significant future direction. The second and third circular statistics were implemented using Astropy package (version 5.1)^23^. The last statistic is state uncertainty which evaluates the mean Euclidean distance between posterior mean and posterior samples of the raw read count prediction: 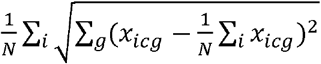 for each cell.

### Cospar model

We computed the Cospar(version 0.1.9)^24^ fate potency score for the above processed multi-fate LARRY dataset, and also selected the subset of the fate potency score for evaluating bi-fate and uni-fate datasets. The following functions achieve this:

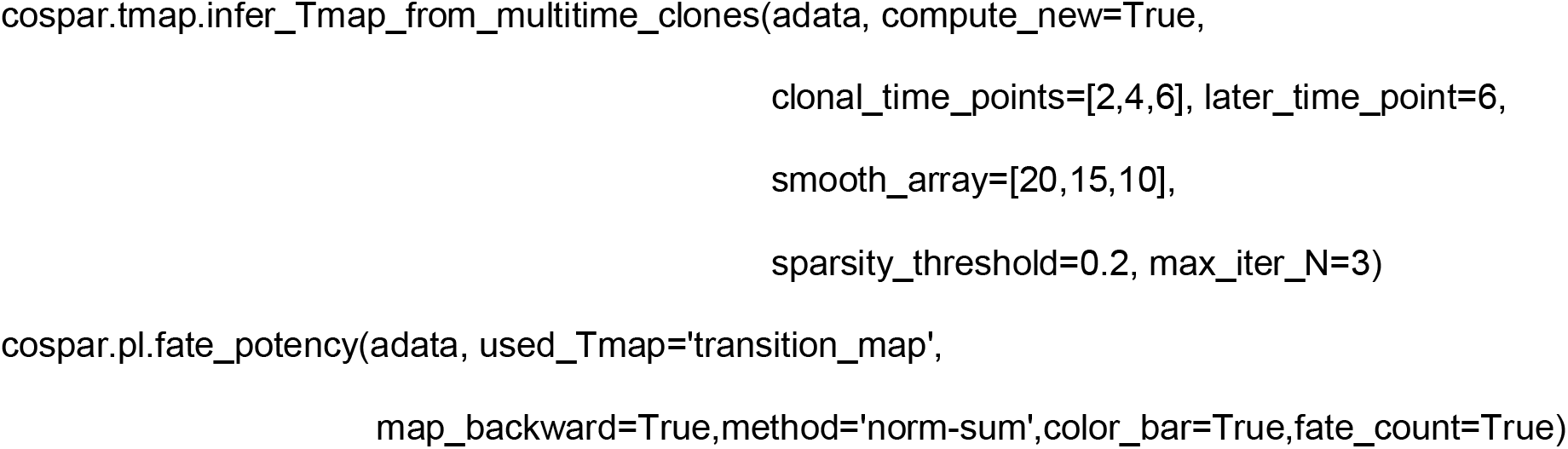

### Cytotrace model

We implemented a faithful Cytotrace Python package based on the R version for benchmarking the Pyro-Velocity and scVelo’s shared (latent) time prediction in the Python environment. This implementation is available in the Pyro-Velocity GitHub repository (https://github.com/pinellolab/pyrovelocity/blob/master/pyrovelocity/cytotrace.py).

### Model evaluation and selection

Our Pyro-Velocity provides two model formulations as described above. Users can compare their predictive performance to decide which of the two models to use for a given dataset. Namely, the mean absolute errors of posterior samples prediction of spliced and unspliced raw read counts *x_icg_* from its observation 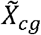 per gene and cell combinations: 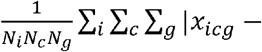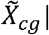. *i* is the posterior sample index, *N_i_* is the posterior sample number, *N_c_* is the cell number, *N_g_* is the gene number. Then it is possible to evaluate the cell fate inference performance of the models based on two metrics: 1. the consistency between the velocity vector field and the *clonal progression vector field* using grid-based cosine similarity; 2. Spearman’s correlation of velocity-based shared (latent) time with Cospar fate potency and the silver standard Cytotrace score. Both metrics are implemented in Scipy.

## Supplementary Figures

**Supplementary Figure 1.**
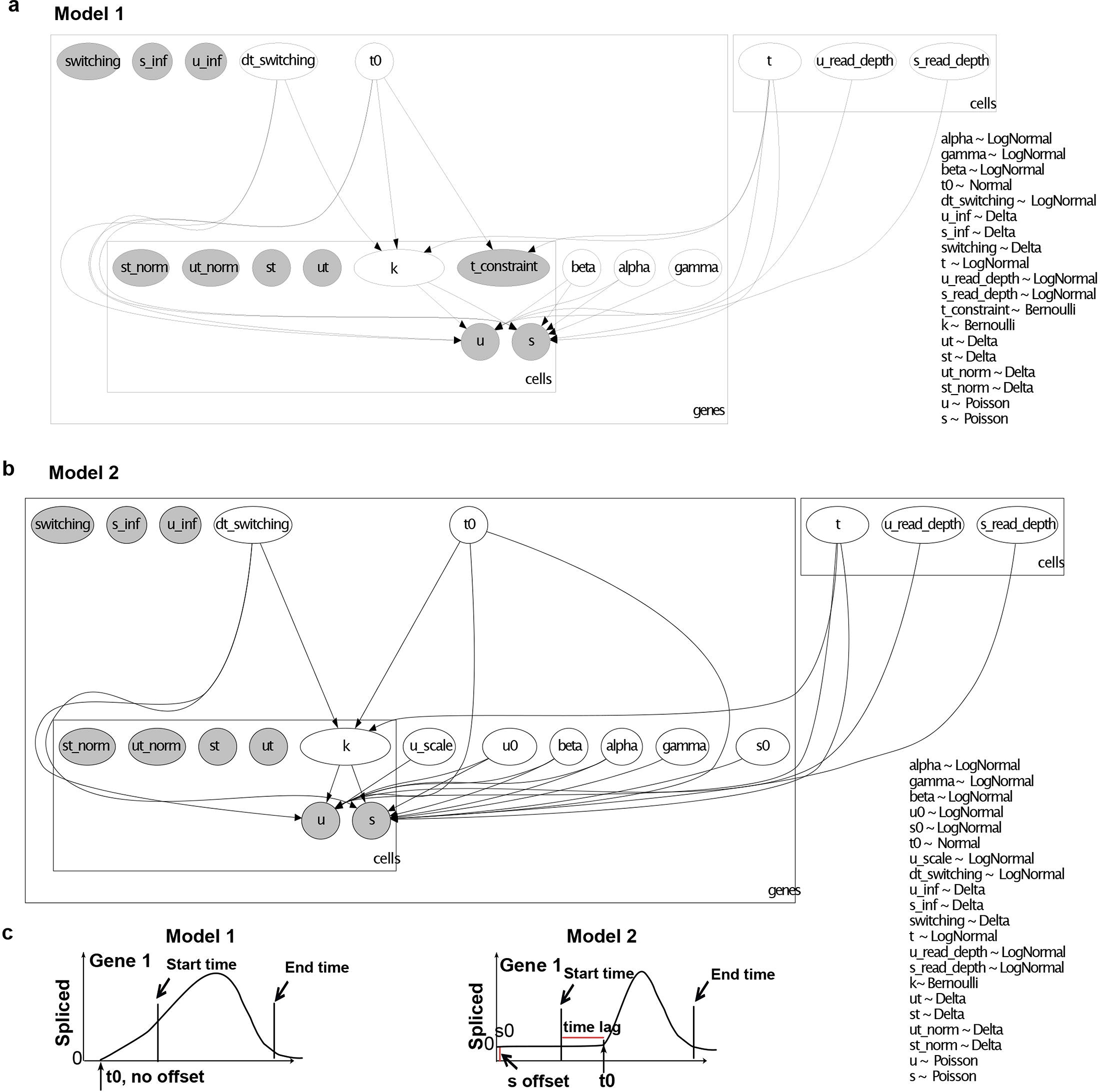
Graphical representations detail the generative process of the Pyro-Velocity models and highlight their differences. **a.** Pyro-Velocity Model 1. Each rectangle represents one data dimension (e.g., gene or cell). The arrows represent the directed hierarchical dependency among random variables. The white circles correspond to the modeled random variables. The probability distributions of these variables are denoted on the right legend. The gray circles correspond to intermediate results computed from random variables. In this model, the kinetics random variables for the cell shared time and gene are modeled with two constraints: 1. the shared time is strictly larger than the gene-specific activation time 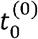, 2. the gene expression in the activate state starts with the initial condition (*u*_0_ = 0, *s*_0_ = 0), as in previous methods^1,4^ (**Methods**). **b.** Pyro-Velocity Model 2. This figure follows the same graphical convention as in a. In this model 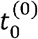 is independent from the shared time, thus allowing a time lag for the gene activation phase. In addition, a positive starting basal gene expression level (*u*_0_, *s*_0_), different than (*u*_0_ = 0, *s*_0_ = 0) is allowed. (Methods). **c.** Schematic illustrating how the constraints proposed in Model 1 (left) and Model 2 (right) are reflected in the modeling of individual gene expression. The x-axis corresponds to the shared time, the y-axis corresponds to the posterior mean of the spliced expression level, the vertical lines with arrows show the start and end time of the observed cell sampling, t0 represents the gene-specific activation time 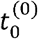, the red lines highlight the differences for the potential gene-specific time lag when 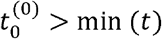, and the initial spliced gene expression. The same differences can also be illustrated for the unspliced gene expression profiles (not shown).

**Supplementary Figure 2.**
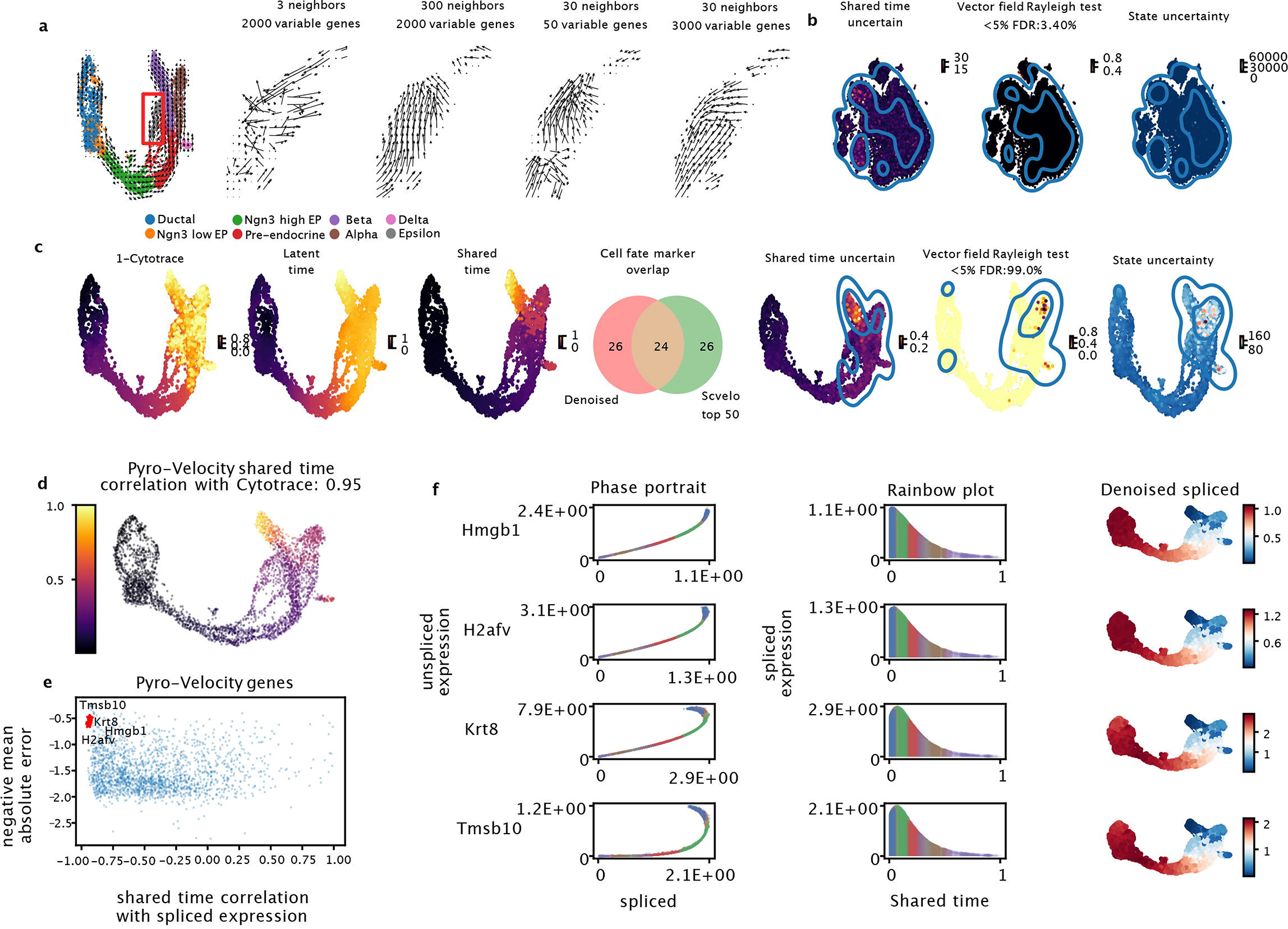
Extended figure of Figure 2 for evaluating Pyro-Velocity (Model 1) and scVelo on the PBMC and pancreas datasets. **a.** Left figure shows the scVelo vector field using the default user options, right figure shows the evaluation of the scVelo vector field using different combinations of user options within Epsilon cells. **b.** Pyro-Velocity additional single cell level uncertainty diagnostic plots on the fully mature PBMC dataset, from left to right: 1. standard deviation of posterior samples of shared time, 2. Rayleigh test of posterior samples of the vector field (the title reports false discovery rate after p-value correction for multiple tests across cells), 3. transcriptional state uncertainty of predicted raw read count per cell. The contour lines show the top 10% of the most uncertain cells. **c.** For the pancreas dataset, the figures show from left to right: UMAP rendering of 1. Cytotrace, 2. scVelo latent time, 3. Posterior mean of Pyro-Velocity shared time averaged across 30 posterior samples, 4. Top 50 markers overlapping between Pyro-Velocity and scVelo (**Methods**, **Supplementary Table 1-2**), 5. The same uncertainty diagnostic plots per cell as in **b. d.** The same plot as **Fig. 2d**. **e.** Selection of the top negatively correlated genes with the cell shared time based on the posterior mean on denoised spliced expression and negative mean absolute error of gene expression prediction (**Methods**). **f.** Phase portraits, rainbow plots, and UMAP rendering of posterior mean of gene expression for the genes selected in e.

**Supplementary Figure 3.**
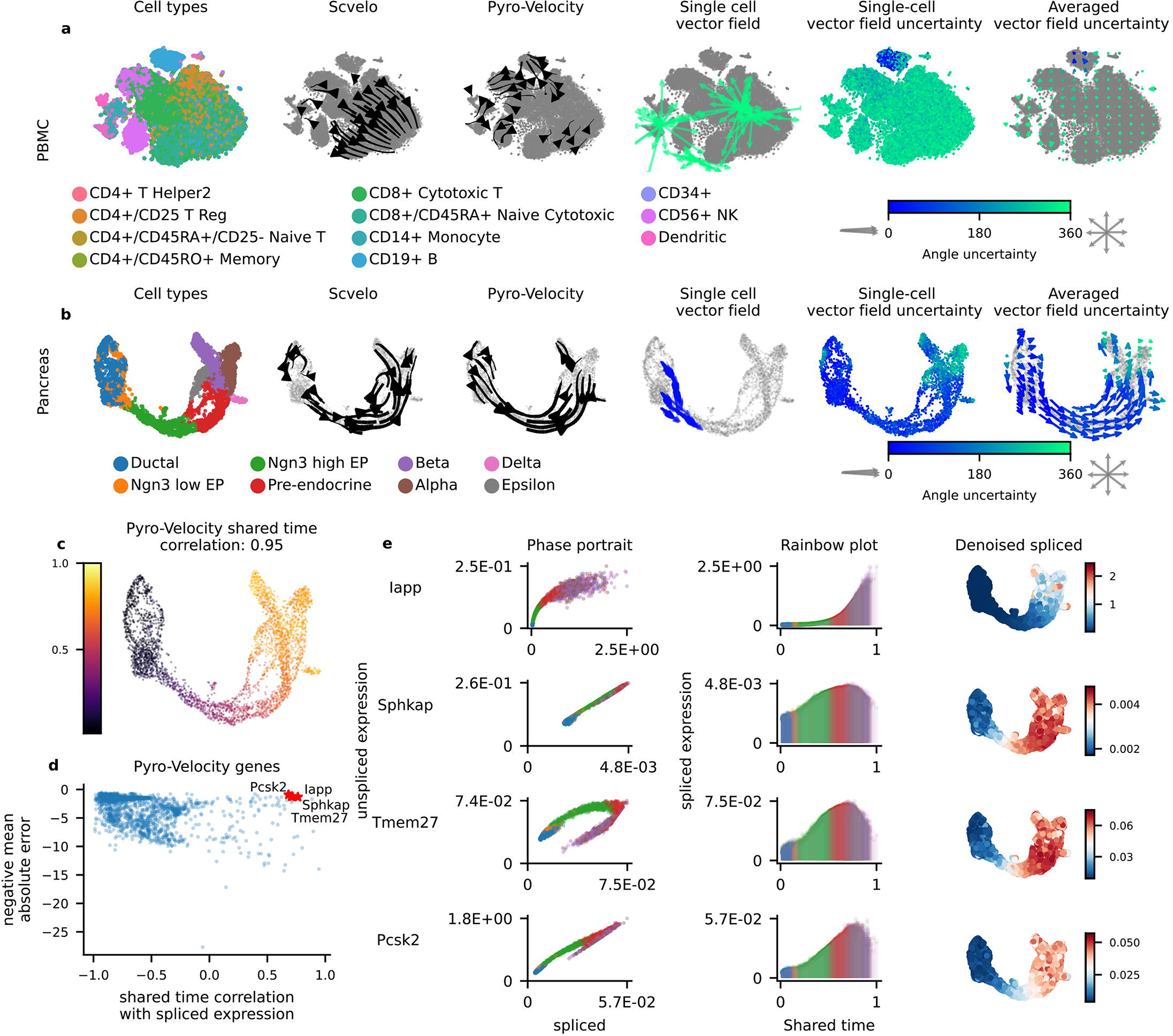
Extended figure of Figure 2, Pyro-Velocity (Model 2) outperforms scVelo on PBMC and pancreas datasets. Same format as in **Fig. 2** and **Supplementary Fig. 2**.

**Supplementary Figure 4.**
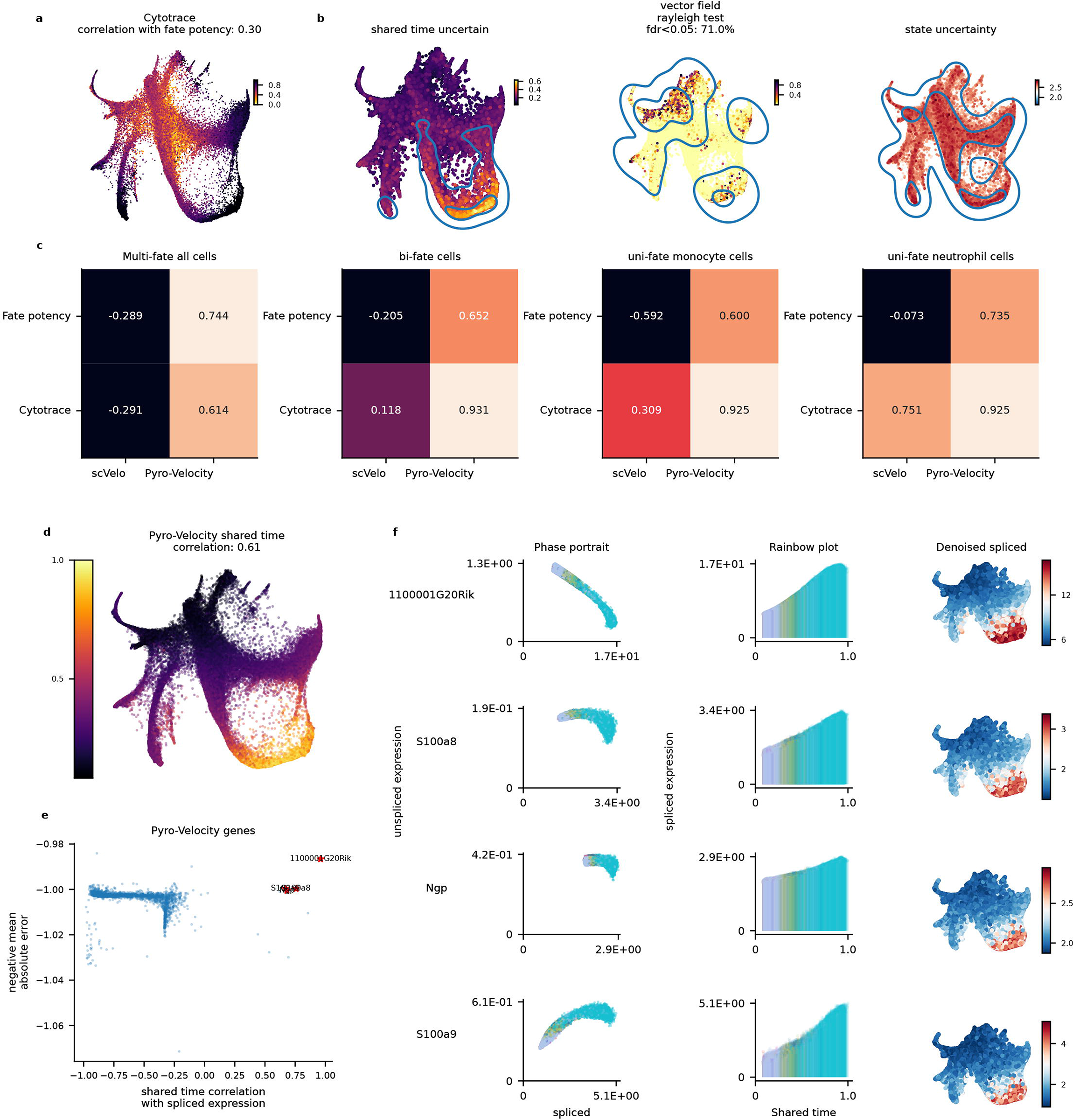
Extended figure of Figure 3, Pyro-velocity (Model 1 for uni-fate data, Model 2 for bi-fate and multi-fate data) outperforms Cytotrace on scRNA-seq datasets with lineage barcoding information (LARRY dataset). **a.** SPRING embedding rendering of Cytotrace scores, the title reports the Spearman’s correlation between the Cytotrace and Cospar fate potency scores, **b.** Pyro-Velocity additional single cell level uncertainty diagnostic plots based on the multi-fate LARRY dataset, from left to right: 1. Standard deviation of posterior samples of shared time per cell, 2. Rayleigh test of posterior samples of vector field per cell (the title reports the false discovery rate after p-value correction for multiple tests across cells), 3. Transcriptional state uncertainty of predicted raw read count per cell. The contour lines show the top 10% of the most uncertain cells (**Methods**)**, c.** Pairwise Spearman’s correlation between Cytotrace/Cospar fate potency scores and velocity methods for the four different benchmarking datasets selected based on the lineage barcoding information from LARRY data. **d.** SPRING embedding rendering of posterior mean of shared time across 30 posterior samples, the title shows the Spearman’s correlation of mean shared time with Cytotrace. **e.** Selection of the top positively correlated genes with the cell shared time based on the posterior mean of spliced expression. **f.** Phase portraits, rainbow plots, and UMAP rendering with splicing gene expression level for the selected genes in e.

**Supplementary Figure 5.**
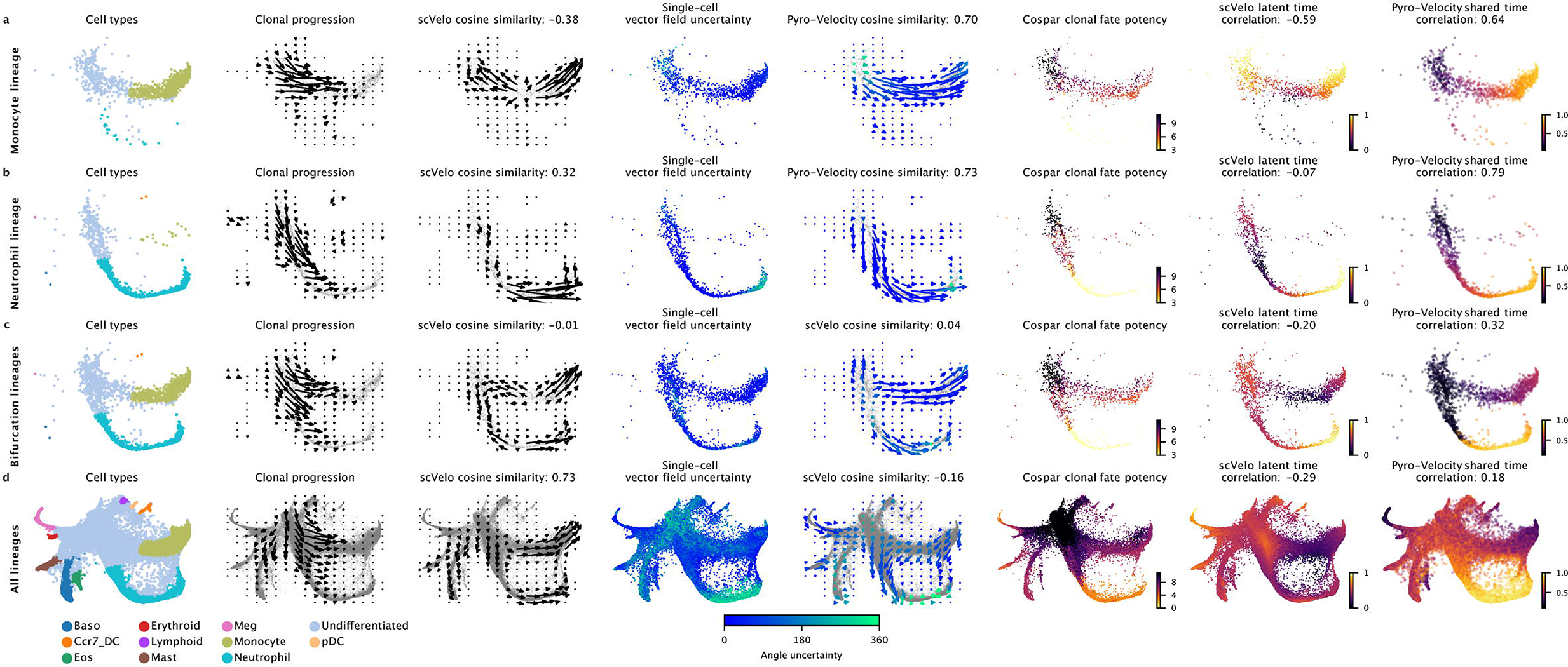
Extended figure of Figure 3 presenting an extended comparison of Pyro-Velocity and scVelo on scRNA-seq datasets with lineage barcoding information (LARRY dataset). **a.** uni-fate Monocyte lineage analysis with Pyro-Velocity Model 2, **b.** uni-fate Neutrophil lineage analysis with Pyro-Velocity Model 2, **c.** Neutrophil and Monocyte bi-fate lineage analysis with Pyro-Velocity Model 1, and **d.** multi-fate lineage analysis with Pyro-Velocity Model 1. The panel descriptions are the same as in Figure 3.

**Supplementary Table 1: Pyro-Velocity Model 1 gene selection comparison with scVelo on the scRNA-seq pancreas dataset.** Sheet 1: Top 50 positively correlated genes with shared time from Pyro-Velocity; Sheet 2: Top 50 positively correlated genes with latent time from scVelo; Sheet 3: Top 50 negatively correlated genes with shared time from Pyro-Velocity; Sheet 4: Top 50 negatively correlated genes with latent time from scVelo.

**Supplementary Table 2: Pyro-Velocity Model 2 gene selection comparison with scVelo on the scRNA-seq pancreas dataset.** Sheet 1: Top 50 positively correlated genes with shared time from Pyro-Velocity; Sheet 2: Top 50 positively correlated genes with latent time from scVelo; Sheet 3: Top 50 negatively correlated genes with shared time from Pyro-Velocity; Sheet 4: Top 50 negatively correlated genes with latent time from scVelo.

## Notes

### Competing Interest Statement

Luca Pinello has financial interests in Edilytics, Inc, Excelsior Genomics, and SeQure Dx, Inc, Luca Pinello interests were reviewed and are managed by Massachusetts General Hospital and Partners HealthCare in accordance with their conflict of interest policies

### Summary of Updates

Updated references

## References

1. La Manno, G. et al. RNA velocity of single cells. Nature 560, 494–498 (2018).

2. Svensson, V. & Pachter, L. RNA Velocity: Molecular Kinetics from Single-Cell RNA-Seq. Molecular cell vol. 72 7–9 (2018).

3. Qiu, X. et al. Mapping transcriptomic vector fields of single cells. Cell 185, 690–711.e45 (2022).

4. Bergen, V., Lange, M., Peidli, S., Wolf, F. A. & Theis, F. J. Generalizing RNA velocity to transient cell states through dynamical modeling. Nat. Biotechnol. 38, 1408–1414 (2020).

5. Bastidas-Ponce, A. et al. Comprehensive single cell mRNA profiling reveals a detailed roadmap for pancreatic endocrinogenesis. Development 146, (2019).

6. Zheng, G. X. Y. et al. Massively parallel digital transcriptional profiling of single cells. Nat. Commun. 8, 14049 (2017).

7. Bergen, V., Soldatov, R. A., Kharchenko, P. V. & Theis, F. J. RNA velocity—current challenges and future perspectives. Mol. Syst. Biol. 17, e10282 (2021).

8. Bingham, Chen & Jankowiak. Pyro: Deep universal probabilistic programming. The Journal of Machine.

9. Li, T. On the Mathematics of RNA Velocity I: Theoretical Analysis. CSIAM Transactions on Applied Mathematics vol. 2 1–55 Preprint at https://doi.org/10.4208/csiam-am.so-2020-0001 (2021).

10. Kucukelbir, Tran, Ranganath & Gelman. Automatic differentiation variational inference. J. Mach. Eng.

11. Gayoso, A. et al. Deep generative modeling of transcriptional dynamics for RNA velocity analysis in single cells. bioRxiv 2022.08.12.503709 (2022) doi:10.1101/2022.08.12.503709.

12. Gorin, G., Fang, M., Chari, T. & Pachter, L. RNA velocity unraveled. PLoS Comput. Biol. 18, e1010492 (2022).

13. Gu, Y., Blaauw, D. & Welch, J. D. Bayesian Inference of RNA Velocity from Multi-Lineage Single-Cell Data. bioRxiv 2022.07.08.499381 (2022) doi:10.1101/2022.07.08.499381.

14. Cui, H., Maan, H., Taylor, M. D. & Wang, B. DeepVelo: Deep Learning extends RNA velocity to multi-lineage systems with cell-specific kinetics. bioRxiv 2022.04.03.486877 (2022) doi:10.1101/2022.04.03.486877.

15. Gao, M., Qiao, C. & Huang, Y. UniTVelo: temporally unified RNA velocity reinforces single-cell trajectory inference. bioRxiv 2022.04.27.489808 (2022) doi:10.1101/2022.04.27.489808.

16. Gulati, G. S. et al. Single-cell transcriptional diversity is a hallmark of developmental potential. Science 367, 405–411 (2020).

17. Millership, S. J. et al. Neuronatin regulates pancreatic β cell insulin content and secretion. J. Clin. Invest. 128, 3369–3381 (2018).

18. Jiang, L. et al. Potential of protein phosphatase inhibitor 1 as biomarker of pancreatic β-cell injury in vitro and in vivo. Diabetes 62, 2683–2688 (2013).

19. Weinreb, C., Rodriguez-Fraticelli, A., Camargo, F. D. & Klein, A. M. Lineage tracing on transcriptional landscapes links state to fate during differentiation. Science 367, (2020).

20. Petukhov, V. et al. dropEst: pipeline for accurate estimation of molecular counts in droplet-based single-cell RNA-seq experiments. Genome Biol. 19, 78 (2018).

21. Melsted, P. et al. Modular, efficient and constant-memory single-cell RNA-seq preprocessing. Nat. Biotechnol. 39, 813–818 (2021).

22. Berens, P. CircStat: A MATLAB Toolbox for Circular Statistics. J. Stat. Softw. 31, 1–21 (2009).

23. Price-Whelan, Sipőcz & Günther. The astropy project: building an open-science project and status of the v2. 0 core package. Astron. J. doi:10.3847/1538-3881/aabc4f/meta.

24. Wang, S.-W., Herriges, M. J., Hurley, K., Kotton, D. N. & Klein, A. M. CoSpar identifies early cell fate biases from single-cell transcriptomic and lineage information. Nat. Biotechnol. 40, 1066–1074 (2022).

